# Face neurons in human visual cortex

**DOI:** 10.1101/2020.10.09.328609

**Authors:** Thomas Decramer, Elsie Premereur, Qi Zhu, Wim Van Paesschen, Johannes van Loon, Wim Vanduffel, Jessica Taubert, Peter Janssen, Tom Theys

## Abstract

The exquisite capacity of primates to detect and recognize faces is crucial for social interactions. Although disentangling the neural basis of human face recognition remains a key goal in neuroscience, direct evidence at the single-neuron level is virtually nonexistent. We recorded from face-selective neurons in human visual cortex, in a region characterized by functional magnetic resonance imaging (fMRI) activations for faces compared to objects (i.e. the occipital face area, OFA). The majority of visually responsive neurons in this fMRI activation showed strong selectivity at short latencies for faces compared to objects. Feature scrambled faces and face-like objects could also drive these neurons, suggesting that the OFA is not tightly-tuned to the visual attributes that typically define whole human faces. These single-cell recordings within the human face processing system provide vital experimental evidence linking previous imaging studies in humans and invasive studies in animal models.

## Introduction

Functional magnetic resonance imaging (fMRI) studies have identified a network of brain regions in occipitotemporal cortex that are activated by faces (Grill-Spector, Knouf, & Kanwisher, 2004; Grill-Spector, Weiner, Kay, & Gomez, 2017; Kanwisher, McDermott, & Chun, 1997; McCarthy, Puce, Gore, & Allison, 1997; Tsao, Moeller, & Freiwald, 2008). The fusiform face area (FFA) (Grill-Spector et al., 2004; Kanwisher et al., 1997) and the occipital face area (OFA) (Gauthier, Skudlarski, Gore, & Anderson, 2000; Pitcher, Walsh, Yovel, & Duchaine, 2007) are considered key components of the human face processing network (Grill-Spector et al., 2017) and yet their exact role in face perception is only partially understood. An important characteristic of these regions, which are often more developed in the right hemisphere (Jonas et al., 2016), is their anatomical consistency across subjects. The OFA is considered the first face-selective region and is thought to respond to the specific features of a face, whereas the FFA, which occupies a higher position in the processing hierarchy, is thought to respond to intact faces and play a role in face recognition (Haxby, Hoffman, & Gobbini, 2000). However, this strict hierarchical view has been challenged and potential interactions between these two nodes have been proposed (Rossion, 2008). For example, studies of prosopagnosia, a disorder of face perception in which patients cannot recognize faces, have linked structural lesions of the right OFA to face recognition (Bouvier & Engel, 2006; Rossion, 2008; Sorger, Goebel, Schiltz, & Rossion, 2007).

Previous intracranial recordings using electrocorticography (ECoG) and depth electrodes (Allison, Puce, Spencer, & McCarthy, 1999; Davidesco et al., 2014; Jacques et al., 2016; Jonas et al., 2016; H. Liu, Agam, Madsen, & Kreiman, 2009; McCarthy, Puce, Belger, & Allison, 1999; Puce, Allison, & McCarthy, 1999; Sato et al., 2014) in face-selective areas have confirmed a preference for faces in these regions; such recordings have improved temporal resolution compared to fMRI, but the signal recorded from macro-contacts still reflects the activity of hundreds of thousands of neurons and can therefore not determine the selectivity of individual neurons. Electrical stimulation over face-selective cortex perturbs face perception (Jonas et al., 2012; Parvizi et al., 2012), thus demonstrating the causal contribution of these areas to face processing.

Face processing in humans may share many similarities with the macaque face processing network, which has been studied extensively at high spatiotemporal resolution using fMRI, single-cell recordings and causal perturbation methods (A. Afraz, Boyden, & DiCarlo, 2015; S. R. Afraz, Kiani, & Esteky, 2006; Chang & Tsao, 2017; Tsao, Freiwald, Tootell, & Livingstone, 2006; Tsao & Livingstone, 2008). Yet, without direct single-cell evidence in humans, the correspondence between the human and the non-human primate remains uncertain.

## Results

We had the unique opportunity to record single-unit (SUA), multi-unit (MUA) and local field potential (LFP) activity in an fMRI-defined, face-selective region in human occipitotemporal cortex (Figure 1A). Based on the CT-MRI coregistration, the microelectrode array (MNI 55, −71, 1; Talairach 56, −70, 5) was located in the anterior superior part of the fMRI activation [faces compared to objects] in occipitotemporal cortex (peak voxel MNI 48, −78, −6) consistent with the OFA (Pitcher, Charles, Devlin, Walsh, & Duchaine, 2009; Pitcher, Walsh, & Duchaine, 2011). This face-selective fMRI activation was part of a larger object-selective activation (contrast intact images of objects compared to scrambled images of objects (Decramer et al., 2019)), which is usually referred to as the lateral occipital complex (LOC). The signal quality of the extracellular recordings was remarkably high, since we detected single-unit activity on multiple electrodes per day (14 to 42 electrodes) with large and easily discriminable waveforms, as illustrated by the example waveform (signal-to-noise ratio of 3.7) in Figure 1B.

**Figure 1.**
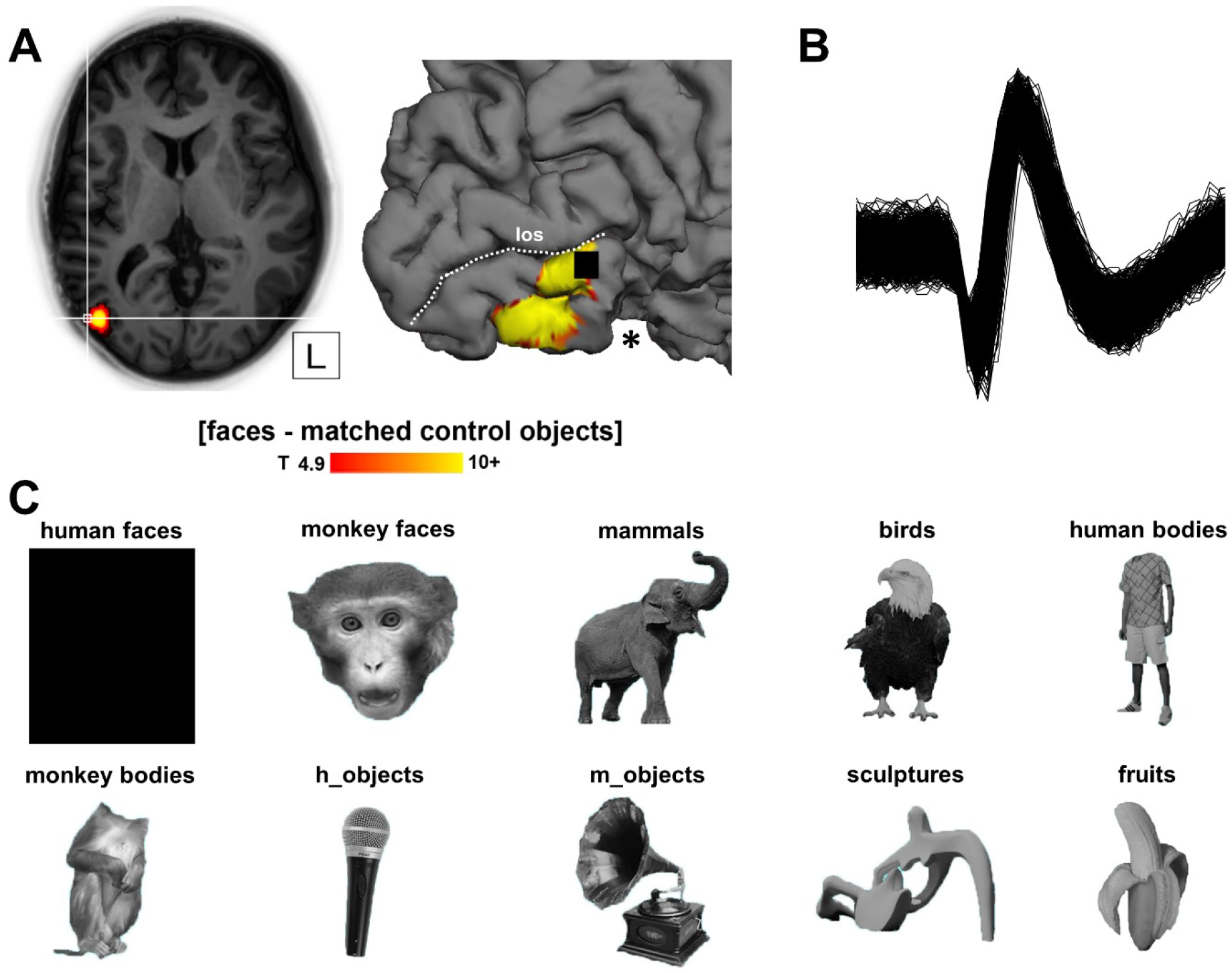
(A) Position of the microelectrode array in relation to fMRI activations, t > 4.9 (p<0.05, FWE corrected for multiple comparisons). The array (black square) is positioned at the border of the fMRI activation below the lateral occipital sulcus (los) in the inferior occipital gyrus of the right hemisphere. The patient underwent a previous resection of inferior occipitotemporal cortex indicated with *, see methods. An overview of the ventral face-selective regions is shown in Figure S1. (B) Example waveform illustrating a high signal to noise ratio. (C) Example stimulus for each image category.

### Category-selective responses in human visual cortex

We recorded SUA (N=67), MUA (N=96) and LFP (high-gamma, N=198) responses to images of different categories (human faces, monkey faces, human bodies, monkey bodies, objects, sculptures and fruits; Figure 1C), as in (Popivanov, Jastorff, Vanduffel, & Vogels, 2014). Many of these visually responsive recording sites showed category-selectivity for faces vs objects (permutation test, p < 0.05; SUA: 36/67, 54%; mean d’ SUA: 0.66; MUA: 57/96, 59%; mean d’ MUA: 0.90; LFP: 147/198, 74%). The example SUA in Figure 2A responded strongly and at short latencies to images of human and monkey faces, but almost did not respond to any of the other categories.

**Figure 2.**
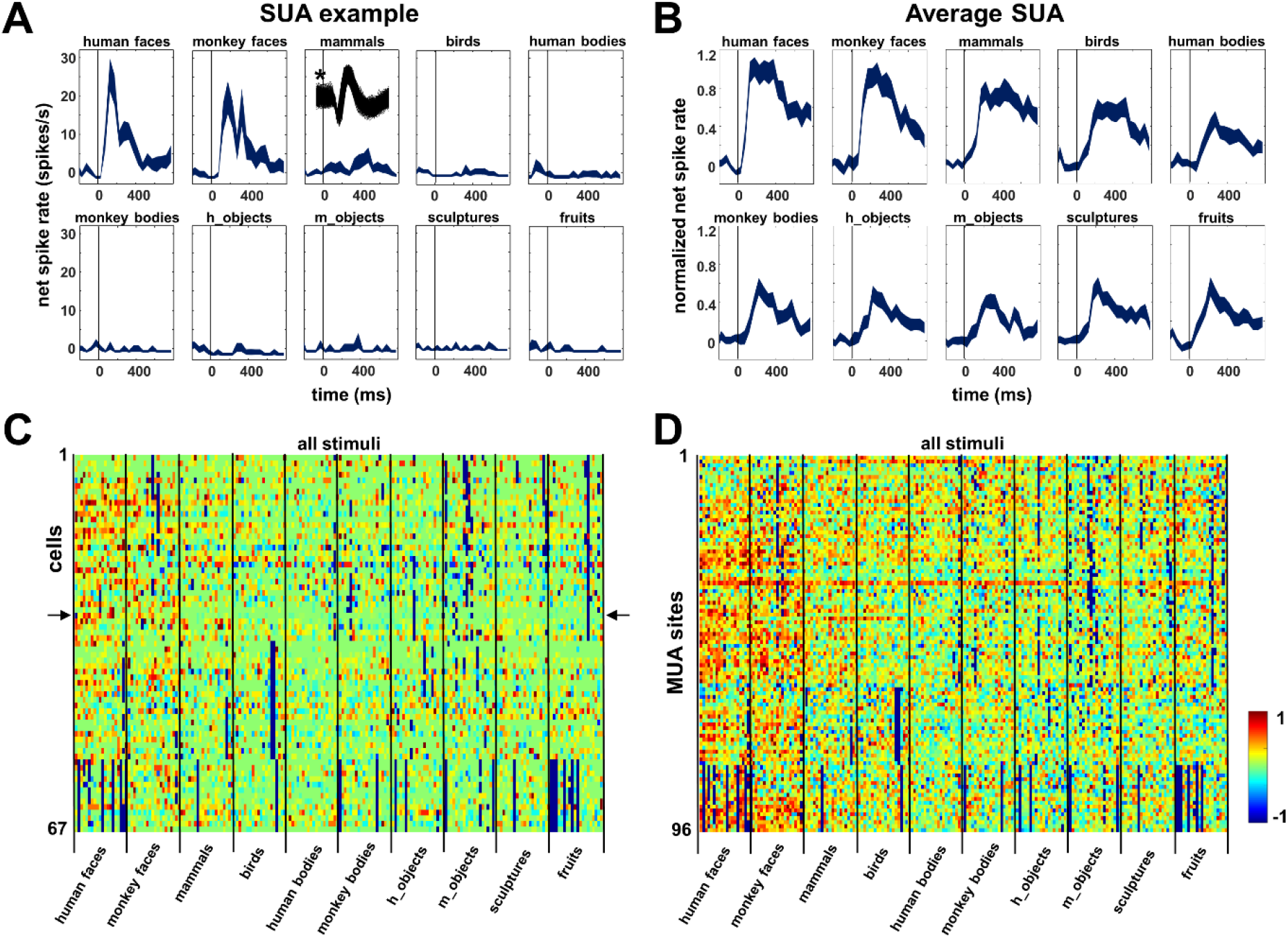
Category selectivity. (A) Example SUA: selective responses to faces (both human and monkey) with almost no response to the other categories; the waveform of the neuron is shown as an inset (*). (B) Average SUA over all visually responsive units (N=67). (C) Net normalized response illustrating between – and within-category selectivity of all visually responsive SUA (N=66), the example SUA is depicted with arrows. (D) Net normalized responses of all visually responsive MUA sites (N=96).

The average response of all visually responsive SUA to faces (N= 67; Figure 2B) emerged very shortly after stimulus onset (around 70 ms; median response latency of 110 ms for SUA, 70 ms for MUA and 79 ms for high-gamma LFP), whereas the average response to objects and bodies was weaker and later (around 120 ms). Figure 2C illustrates the between – and within-category selectivity of all visually responsive SUA (N= 67) by plotting the net normalized responses to every stimulus in the test. The example neuron from Figure 2A responded exclusively to faces (see arrows in Figure 2C). It is clear that both the human and the monkey face categories evoked the strongest responses, but individual stimuli from the other categories could also activate these neurons. The average response to mammals and birds (which also contained faces) was lower than to face stimuli, and the other stimulus categories evoked even weaker responses. Figure 2D illustrates the between – and within-category selectivity of all responsive MUA sites (N =96), showing similar results. Overall, consistent with the fMRI localizer, we recorded in a relatively homogeneous face-selective patch of object-selective visual cortex.

Our category test contained images of faces, objects and bodies. To illustrate the neuronal selectivity for faces compared to bodies across the array, we calculated a d’ index comparing the MUA responses to faces and bodies (Figure 3A). Many electrodes exhibited strong selectivity (d’ > 1; N=15) for faces compared to bodies (Figure 3B), but we also observed a small number of electrodes with stronger responses to bodies than to faces (2 MUA and 2 SUA, blue squares in Figure 3A and example recording site in Figure 3C). These body-selective sites typically responded to faces (and in the case of the example site also to objects), but much stronger to images of human bodies, and were located merely 1 to 3 mm from the face-selective sites. Therefore, although dominated by face-selective responses, the human OFA also contains body-selective sites in close proximity to clusters of neurons preferring faces.

**Figure 3.**
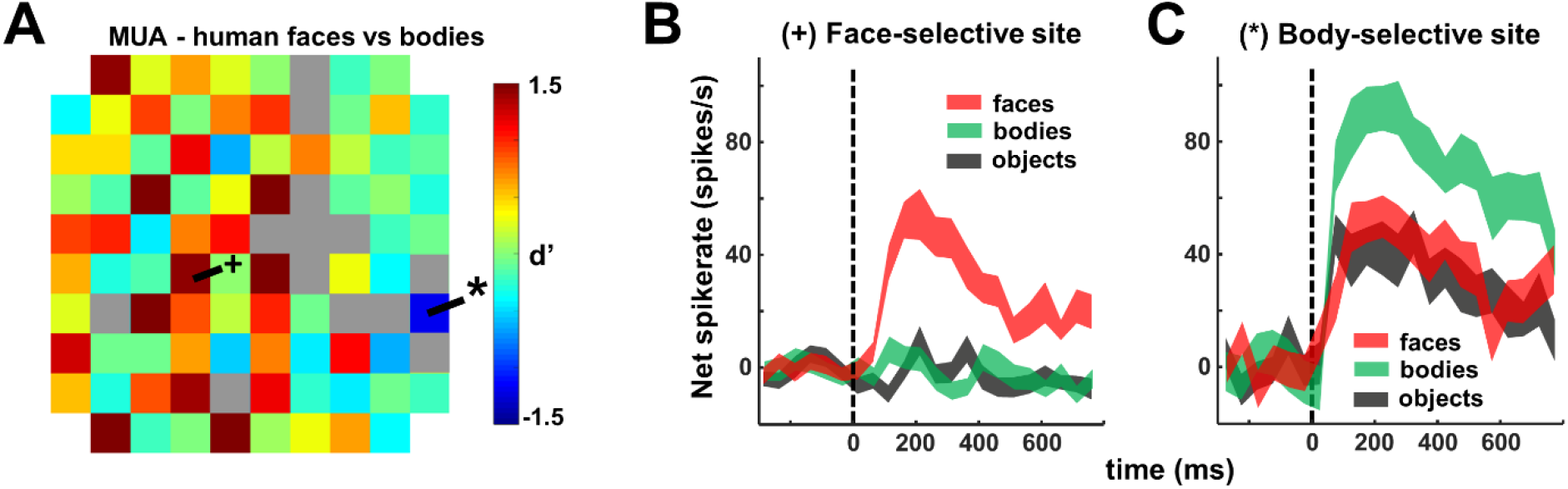
Face versus body selectivity. (A) MUA d’ values across the array, illustrating a face (+) and a body (*) selective site. (B) Corresponding MUA response profiles of face selective site. (C) Corresponding MUA response profile of body-selective site.

Next, we wanted to determine to what extent face cells (defined as faces>objects from the category experiment; SUA N=10, MUA N=18) in the OFA differentiate between human faces. For this test, we presented images of famous faces (actors, politicians, N=10), faces of people familiar to the patient (N=2, her mother and the neurosurgeon) and a set of cartoon faces (N=6) (Figure 4A). Overall, most of the face cells (SUA N=7/10, MUA N=11/18) showed differential responses within the faces category (ANOVA p < 0.05, Figure 4B; 6/10 SUA and 6/18 MUA sites were significantly selective for the human faces, excluding the cartoon faces). To quantify the width of the selectivity, we calculated a selectivity index defined as S_width_ = (n - ∑r_i_ / max) / (n-1) (Rainer, Asaad, & Miller, 1998), which is equal to 1 if the neuron only fires to a single stimulus in the test, and 0 if the neuron fires equally to every stimulus. In our face test, the S_width_ averaged 0.44 (ranging from 0.73 to 0.26) for MUA and 0.75 (ranging from 1.0 to 0.58) for SUA, indicating that although the face cells in the OFA respond to a wide range of faces, a considerable amount of within-category selectivity was present. The average MUA and SUA (Figure 4C) responses in the test also indicate that we measured significant responses to every face stimulus in the test, without apparent preference for famous, familiar or cartoon faces. Using a famous face, we also performed a receptive field (RF) mapping of these neurons. The RFs were large and bilateral with a contralateral preference (Figure S2).

**Figure 4.**
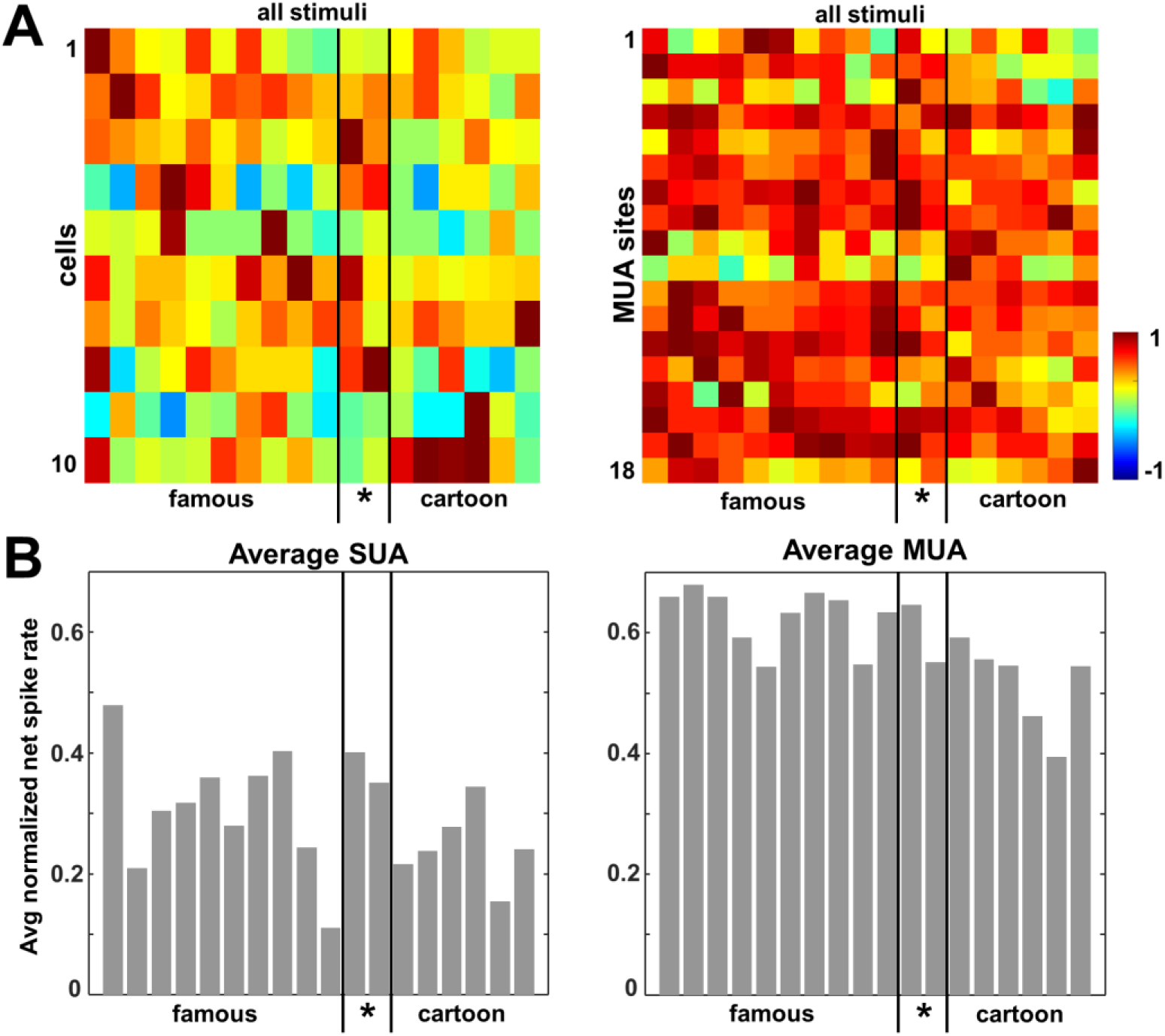
Face selectivity. (A) Normalized SUA (left) and MUA (right) response for each site and stimulus (famous persons, familiar faces (*) and cartoon faces). (B) Average normalized net response for all sites per stimulus.

The results we obtained using famous faces demonstrate that the face cells we recorded in human visual cortex were broadly tuned to face stimuli. Do these neurons respond to discrete facial features or whole intact faces? To address this question, we investigated how these neurons responded to feature scrambled faces, in which the position of the face elements (eyes, nose, lips and mouth) was altered. The example neuron fired to the intact face (Figure 5A first column), while the feature scrambled face at the same location evoked a virtually identical response (Figure 5A, second column, t-test, p = 0.102), indicating that the spatial configuration of the face elements was not crucial for this neuron. However, stimuli in which the face elements were further dispersed in the visual field (Figure 5A, third and fourth column) evoked significantly less response (t-test, p =0.032 & p <0.001 respectively) compared to the intact face. Overall, most single neurons (4/6, 67%) and MUA sites (14/17, 82%) did not respond more to their preferred intact face than to the corresponding feature scrambled face in which the face elements occupied the same position in the visual field. We observed a similar phenomenon at the level of the LFP (Figure 5B). The normalized net MUA responses to intact vs. feature scrambled faces for each MUA site are depicted in two scatter plots (Figure 5C), illustrating that most sites also responded to feature scrambled faces, even when the face elements are spread further apart. These data demonstrate that face neurons in the human OFA are not only driven by intact faces with their features in the canonical configuration, but also other spatial recombinations of facial features. Since the configuration of facial features, especially distances between features, is thought to carry diagnostic information essential for face recognition (Chang & Tsao, 2017; Issa & DiCarlo, 2012), this observation is consistent with a role for the OFA in face detection.

**Figure 5.**
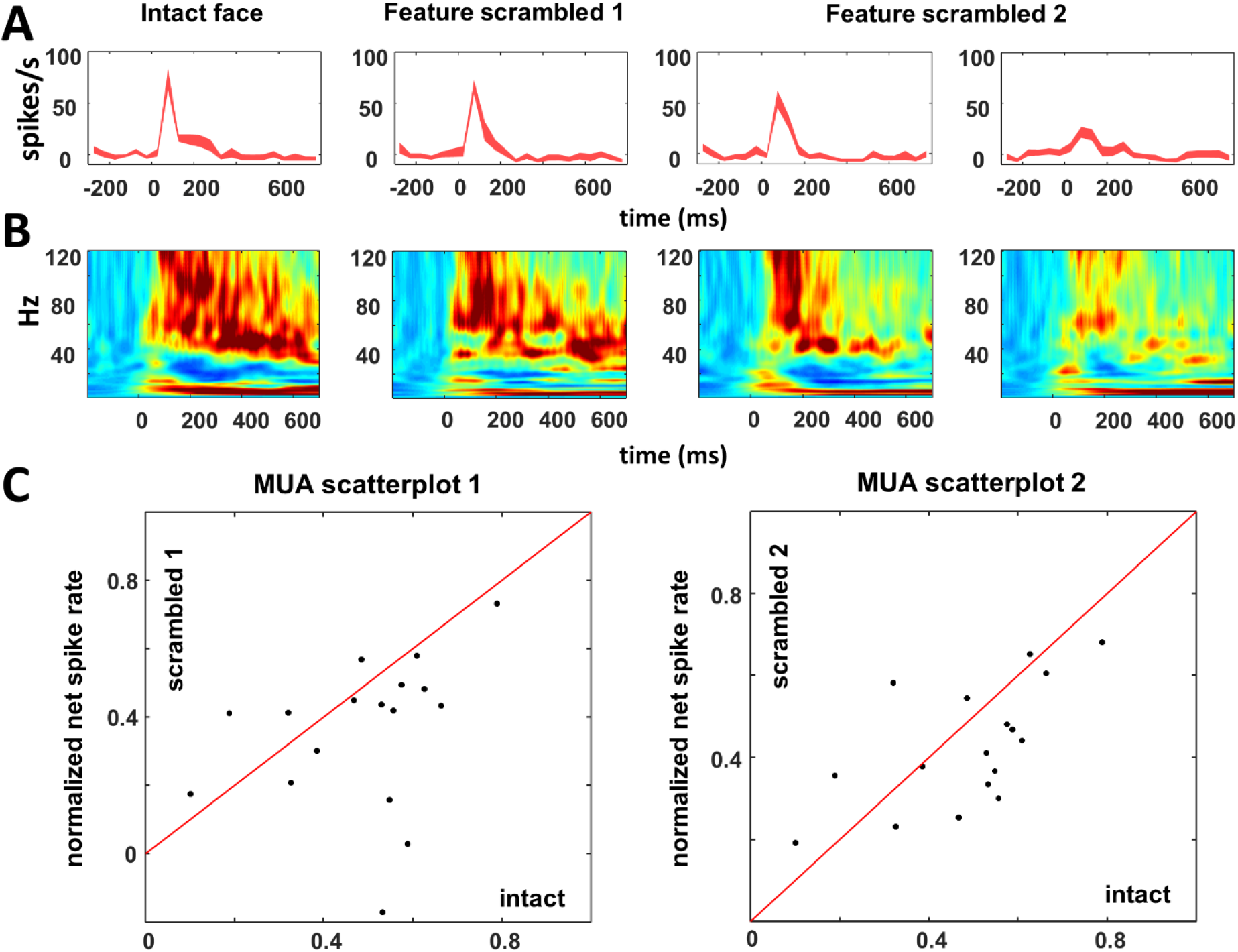
Responses to feature scrambled faces. (A) Example SUA. Feature scrambled 1 refers to the stimuli in which the face elements occupied the same position in the visual field, feature scrambled 2 refers to stimuli in which the face elements were further dispersed in the visual field. (B) Average LFP spectrum over all visually responsive sites (N=39). (D) MUA scatter plot for intact vs feature scrambled faces at same location (left) and for intact vs feature scrambled faces with the face elements further apart (right).

### Face pareidolia

To further interrogate tuning of facial features at the single-cell level, we capitalized on the phenomenon of face pareidolia (Figure 6A), the compelling illusion of perceiving illusory facial features in inanimate objects, which is experienced by both humans (J. Liu et al., 2014; Nihei, Minami, & Nakauchi, 2018; Wardle, Seymour, & Taubert, 2017) and macaque monkeys (Taubert, Wardle, Flessert, Leopold, & Ungerleider, 2017). This phenomenon is relevant because, unlike real faces, examples of face pareidolia (hereafter referred to as ‘face-like objects’) have facial features that are highly variable in terms of their visual attributes. For example, unlike the features of faces, which tend to be skin-colored and round in shape, face-like objects have features that vary in both color and shape. In the fMRI activation [*faces* – *objects*], the percent signal change evoked by face-like objects was intermediate between that evoked by faces and by objects (Figure 6C). We recorded neural responses to images of faces, face-like objects and control objects, which were matched in object identity to the face-like objects (Taubert et al., 2017) (Figure 6A).

**Figure 6.**
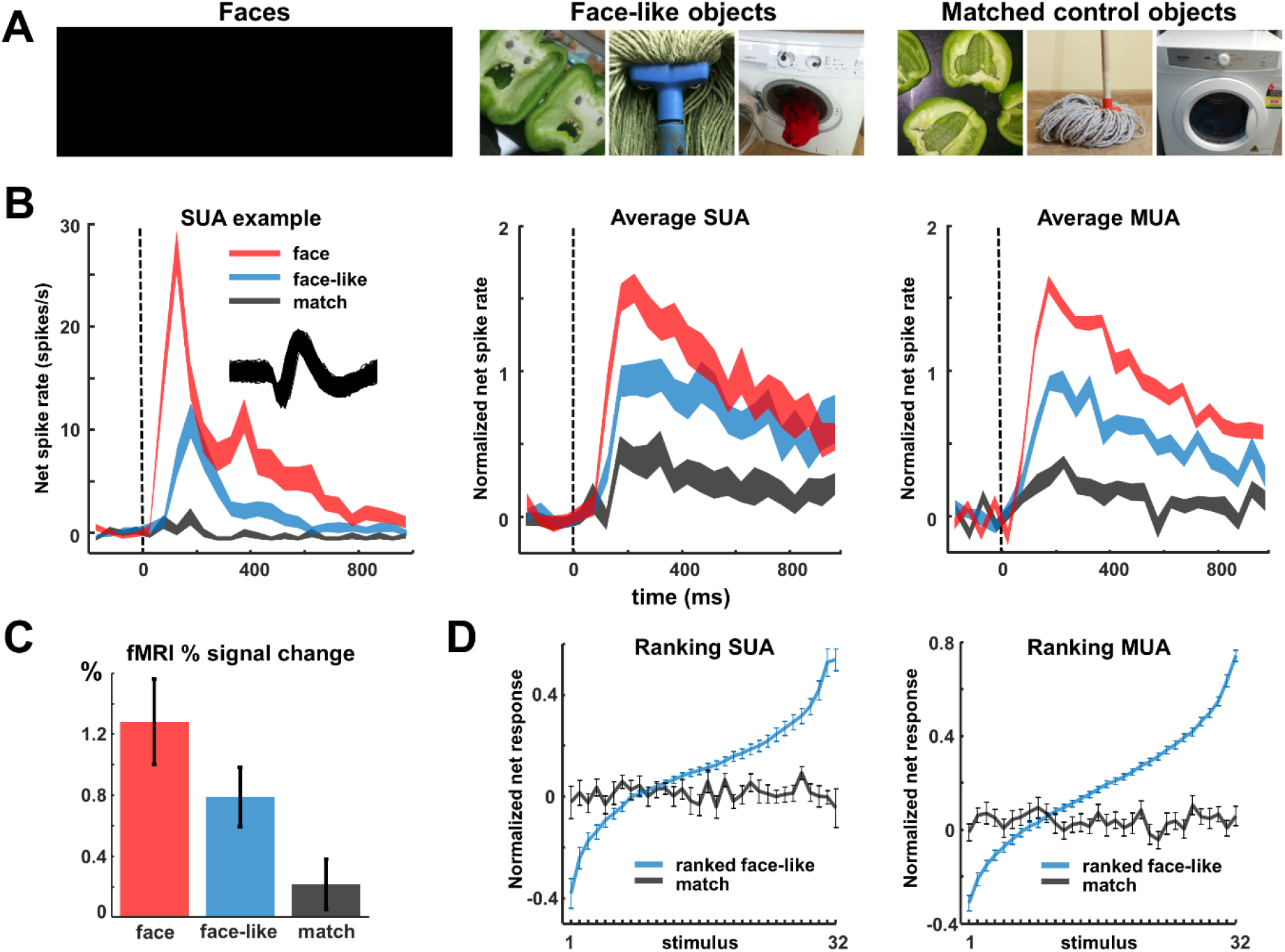
Face pareidolia. (A) Example stimuli of faces, face-like objects and matched control objects. (B) SUA example illustrating the intermediate response to face pareidolia. Average SUA (N=44) and MUA (N=50) over all visually responsive sites. (C) fMRI percent signal change at the right OFA 6w after removal of the microelectrode array. (D) Average ranking for the 32 face-like stimuli for both SUA and MUA. For the matched control objects, the ranking is lost.

The example neuron in Figure 6B responded strongly to faces and weakly to control objects. The response to face-like objects was distinctive, since it was greater than the response elicited by non-face objects, yet weaker than the response to real faces. For this example neuron recorded in human visual cortex, the average response to face-like objects (14 spikes/s) was less than half of that to faces (31 spikes/s), and significantly delayed compared to the responses to real faces (peak response at 170ms for face-like objects vs 90ms for faces). In total, we recorded the activity of 44 visually responsive single neurons to faces, face-like objects and matched control stimuli, the majority of which (26/44, 59%) were selective for faces compared to objects. Averaged over all visually responsive single-units, the normalized response to faces was significantly stronger than that to control objects, whereas face-like objects elicited an intermediate response (Figure 6B, middle panel). This pattern was even more pronounced for the multi-unit activity (N=50, 72% were selective for contrast face > matched control object, Figure 6B right panel) and for the average high-gamma (80-120Hz) band responses of the LFP (N=111), almost all of which (109 or 98%) were selective for the contrast face > object. Note that the proportion of selective sites was significantly higher for the high-gamma responses compared to the SUA and MUA data (z-test for proportions, p < 0.05). The median selectivity latency (faces > control objects) was 130 ms for SUA, 110 ms for MUA and 108 ms for LFP (Supplementary Figure S3). The preference for real faces within our neuronal population was very robust, since only three single neurons and no MUA or LFP site preferred face-like stimuli over faces (at p < 0.05, Supplementary Figure S4). Moreover, no SUA, MUA or LFP site preferred control objects over faces (at p < 0.05).

To assess the relative importance of face and object features, we ranked the face-like objects based on their average normalized SUA responses for each cell separately, calculated the average response to the ranked stimuli across channels (Figure 6D), and compared the responses in this ranking to the corresponding matched control objects (i.e. stimuli with the same object identity). This procedure was repeated for MUA (for ranking of face stimuli, see Figure S5). The slopes of the linear fits to these curves reveal the neuronal selectivity for face-like objects and whether this selectivity was preserved for control objects. Figure 6D clearly shows that the selectivity for face-like stimuli was considerable: only 19% (for SUA), 22% (for MUA) and 31% (for high gamma) of the stimuli evoked responses above 50% of the maximum response. However, no systematic preference was present for the corresponding control objects, since the slopes of the regression lines were not significantly different from zero (see Table S1), indicating that the presence of face-like features is critical to drive neurons in this face-selective region.

## Discussion

With the exception of one study showing spiking activity near the human FFA (Axelrod et al., 2019), no study has reported detailed single-neuron responses in an fMRI-defined face-selective region in human visual cortex. In a manner analogous to face cells in the temporal cortex of the macaque monkey, individual human occipitotemporal neurons showed strong and early selectivity for faces, consistent with fMRI activations. Feature scrambled faces were often effective in driving the neuronal responses, and face-like objects also elicited reliable responses from this face-selective region. Thus, neurons at this level of the visual hierarchy are broadly tuned to the features of a face, independent of their spatial configuration and their low level visual attributes, such as color.

The category selectivity we observed in the OFA was less pronounced than in some of the macaque face patches (Tsao et al., 2006). Approximately half of the visually responsive SUA and MUA sites were not category-selective, and we even observed a few body-selective sites on the same 4 by 4 mm electrode array. These observations could at least partially be explained by the fact that the electrode array was located at the edge of the fMRI activation [faces – objects]. A previous study in the macaque middle face patch reported a gradient in the number of face-preferring sites from the center towards the periphery (Aparicio, Issa, & DiCarlo, 2016), see also (Bell et al., 2011).

A number of observations suggest that the human OFA represents a relatively early stage in the face-processing hierarchy. The face cells also responded to feature scrambled faces, their response latencies were short (as early as 70 ms), the tuning within the face category was relatively broad and face-like stimuli also activated the neurons. On the other hand, RFs were large and bilateral, consistent with earlier fMRI studies of the OFA (Hemond, Kanwisher, & Op de Beeck, 2007), and we observed considerable selectivity within the face category compared to the most studied face patch in the macaque monkey (the middle lateral (ML) face patch, (Tsao et al., 2006)). A possible monkey homologue for the OFA is the posterior lateral (PL) face patch (Tsao et al., 2008). Single neuron recordings in monkeys have shown that the presence of an eye with a contour can drive neurons in PL like real faces (Issa & DiCarlo, 2012). In the absence of single-unit recording data in other face-selective regions in human visual cortex, however, it is difficult to draw strong conclusions about possible homologies with the macaque face patches (see also (Rossion & Taubert, 2019)).

The latency of the face responses we report here (70 ms) was shorter than the latency of the responses to images of objects in our previous study (120 ms, (Decramer et al., 2019)), which was in the same area. This apparent discrepancy is most likely related to the fact that the face stimuli evoked stronger responses compared to images of objects, which has a marked effect on the latency estimation especially with a limited number of trials. Note also that SUA, MUA and high-gamma responses were consistent with the pattern of fMRI responses to faces, face-like stimuli and objects, although the number of face-selective sites was much higher at the level of the high-gamma responses compared to SUA. Thus, our results also represent a unique validation of an extensive body of research using fMRI to study face processing in human subjects (Grill-Spector et al., 2004; Grill-Spector et al., 2017; Kanwisher et al., 1997).

Single neurons selective for faces have been reported in the medial temporal lobe (MTL) of epilepsy patients (Fried, MacDonald, & Wilson, 1997). These face cells respond with a much longer latency of around 300ms (Kreiman, Koch, & Fried, 2000), are multimodal (i.e. also respond to written (Quiroga, Reddy, Kreiman, Koch, & Fried, 2005) and spoken names (Quian Quiroga, Kraskov, Koch, & Fried, 2009)), and frequently signal face identity. These medial temporal regions occupy a much higher level in the face processing network (Tsao et al., 2008), explaining their late responses.

A limitation of our study is the fact that this patient previously underwent a resection of a cystic lesion in the ventral occipitotemporal cortex. The right FFA might have been damaged during previous surgery, or the OFA-FFA network might have developed differently due to the presence of the lesion. Moreover, intracranial recordings with depth electrodes revealed that the patient had mesial temporal lobe epilepsy, which can affect the face processing network (Riley, Fling, Cramer, & Lin, 2015). The patient did not suffer from prosopagnosia, a condition which has been linked to the right OFA (Pitcher et al., 2011; Rossion, 2008).

Although human intracortical recordings remain scarce, they provide an extraordinary opportunity to study the human brain with unparalleled spatiotemporal resolution. Neural recordings in fMRI-defined patches in human cortex are critical for validating human fMRI data and provide a novel way to bridge the gap between human imaging and invasive studies in animal models.

## Methods

Ethical approval was obtained for microelectrode recordings with the Utah array in patients with epilepsy (study number s53126). Study protocol s53126 was approved by the local ethical committee (Ethische commissie onderzoek UZ / KU Leuven) and was conducted in compliance with the principles of the Declaration of Helsinki (2008), the principles of good clinical practice (GCP), and in accordance with all applicable regulatory requirements. Strict adherence to all imposed safety measures, including case report forms (CRFs) and in detail reports on (serious) adverse events ([S]AEs), was required. All human data were encrypted and stored at the University Hospitals Leuven.

In patients with intractable epilepsy, invasive intracranial recording studies aimed to identify the EZ for subsequent removal; microelectrodes (Utah array) were additionally implanted to study the microscale dynamics of the epileptic network in the presurgical evaluation for research purposes. No additional incisions were made for the purpose of the study. The Utah array was placed on the convexity of the brain at the target location of—and in combination with—clinical electrodes, analogous to previous studies using microelectrode arrays, such as (Martinet et al., 2017; Smith et al., 2016; Truccolo et al., 2011). Target locations for clinical electrodes were determined by the epileptologist and were based on preoperative electroclinical and advanced imaging investigations (including MRI, PET, SPECT/SISCOM). The location of the Utah array was always at the site of the presumed epileptogenic zone (PEZ) and away from eloquent brain areas, as determined by preoperative task-based motor and language fMRI.

During surgery, a microelectrode array was implanted in the right occipitotemporal cortex, in an area of cortex with a high probability of resection provided that the presumed ictal onset zone was confirmed after invasive intracranial recordings. The patient had undergone a surgical resection of a cystic lesion (WHO grade I ganglioglioma; MNI 28 to 46, −42 to −62, −5 to −27 mm) in the right occipitotemporal region 7 y earlier, with late seizure recurrence after initial seizure freedom. The patient was on brivaracetam 50 mg twice daily and perampanel 8 mg once daily. Intracranial recordings revealed an anterior temporal EZ and the patient underwent a temporal lobectomy including the mesial temporal structures several months later.

Written informed consent was obtained before the start of the study. Possible additional risks of microelectrode placement, such as superficial hemorrhage and infection were discussed on 2 different occasions (once at the outpatient clinic and again before surgery). The Utah array was used because this represents the only FDA-approved microelectrode array, capable of studying the microscale dynamics of the epileptic network at the single-unit level, and because these arrays have been safely used in previous studies in both epilepsy patients (Martinet et al., 2017; Smith et al., 2016; Truccolo et al., 2011; Weiss et al., 2013) as well as in patients undergoing BMI implants (Ajiboye et al., 2017; Collinger et al., 2013; Davis et al., 2016; Flesher et al., 2016; Hochberg et al., 2012; Hochberg et al., 2006; Simeral, Kim, Black, Donoghue, & Hochberg, 2011). Classical clinical electrodes have large contact surfaces and record from a large number of neurons (10^4^–10^5^) and thus are not capable of studying microscale dynamics. As these microelectrode arrays are not CE marked, local regulatory approval (FAGG) was additionally obtained. Previous histological studies show that microelectrode arrays cause focal tissue damage at the thin electrode trajectories upon insertion (Fernandez et al., 2014; House, MacDonald, Tresco, & Normann, 2006). Postoperative MRI several weeks after electrode removal did not show any signal changes at the implantation site, indicating that there were no larger cortical lesions associated with array implantation.

To minimize trauma and risk of hemorrhage due to microelectrode insertion, we performed under-insertion of the array with a low insertion pressure (15 psi). This implies that the pressurized inserter did not insert the electrodes over their full length, ensuring that the electrode pad on which the electrodes are bonded does not impact the brain. The risk of infection was minimized by using a custom percutaneous connector, with a small wire tunneled through the skin, in analogy to the classic clinical electrodes.

### Functional magnetic resonance imaging (fMRI)

fMRI was performed 3 months after array extraction on a 3T scanner (Achieva dstream, Philips Medical Systems, Best, The Netherlands) in one session of 60 minutes. Functional images were acquired using gradient-echoplanar imaging with the following parameters: 52 horizontal slices (2 mm slice thickness; 0.2 mm gap; multiband acquisition), repetition time (TR) 2 s, time of echo (TE): 30 ms, flip angle 90°, 112 * 112 matrix with 2 x 2 mm in plane resolution, and sensitivity enhancing (SENSE) reduction factor of 2. Stimuli were projected with a liquid crystal display projector (Barco Reality 6400i, 1,024 x 768 pixels, 60-Hz refresh rate) onto a translucent screen positioned in the bore of the magnet (57 cm distance). The patient viewed the stimuli through a mirror tilted at 45° and attached to the head coil. Stimuli were presented for 1000 ms (interstimulus interval, 0 ms) on a grey background, with a stimulus size of 8° and a red fixation square of 0.4°. Two runs of a block-design task using stimuli consisting of human faces (24s), face pareidolia (24s), matched control objects (24s) and fixation (12s) with 4 repeats within one run (336s) were performed.

Data analysis was performed using the SPM12 software package (Wellcome Department of Cognitive Neurology, London, UK) running under MATLAB (The Mathworks, Natick, MA). Preprocessing involved realignment of the images followed by co-registration of the anatomical image and the mean functional image. Before further analysis, the functional data were smoothed with an isotropic Gaussian kernel of 5 mm. On Figures 1 and S2, all activations with t > 4.9 (p<0.05, FWE corrected for multiple comparisons) are shown for the contrast [faces – matched control objects]. We calculated the mean percentage signal change (using MarsBaR v0.41.1) for the two runs for each condition vs fixation and verified the main effect of condition by performing a one-way ANOVA.

### Microelectrode recordings

The array was interfaced with a digital headstage (Blackrock Microsystems, UT, USA) connected to a 128-channel neural signal processor (Blackrock Microsystems, UT, USA). Spiking activity was high-pass filtered (750 Hz). A multi-unit detection trigger was set at 95% of the signal’s noise. For LFP recordings, the signal was filtered with a digital low-pass filter of 250 Hz, with a sampling frequency of 1000 Hz. Spike sorting was performed offline (Offline Sorter 4, Plexon, TX, USA). On five channels, we isolated two visually responsive single-units.

### Stimuli and tests

Stimuli were presented in a custom-made stereoscope. Images from two LCD monitors were presented to the right and left eyes with the use of customized mirrors at a viewing distance of 56 cm (1 pixel = 0.028°). The patient was instructed to fixate a small target at the center of the display.

#### Category test

The patient had to discriminate between achromatic photographs of birds (target) and other stimuli (distractors: mammals, fruits, bodies (human-monkey), faces (human-monkey), objects (controlled for human bodies and controlled for monkey bodies, sculptures) displayed in the center of the screen (stimulus size: 7°), for each category there were 20 stimuli (Popivanov et al., 2014). Birds appeared randomly in 10% of trials. All stimuli were presented for 800 ms. We recorded on day 2, 4 and 6 after array implantation and pooled the data over these three sessions.

#### Famous faces test

The patient passively viewed achromatic images of famous persons (n=10), familiar faces (n=2; her mother and the senior neurosurgeon (T.T.)) and cartoon faces (n=6) displayed at the center of the screen (stimulus size: 6°). Stimulus presentation was 1000ms. We recorded on day 6 after array implantation.

#### Receptive field test

To map the RF, a single famous face stimulus (6°) was presented at 25 different positions in the visual field, covering 50 degrees horizontally and 30 degrees vertically, during passive fixation.

#### Feature scrambled faces test

The patient passively viewed achromatic images of faces (stimulus size: 6°) at four different positions around the fixation point. Feature scrambled stimuli with the same contour were presented at the same location and several feature scrambled stimuli with face components over the visual field were presented. Each stimulus had two exemplars (male and female). We recorded on day 8 after array implantation.

#### Face pareidolia test

Stimuli consisted of 32 real faces, 32 face-like objects and 32 matched control objects. Two handheld buttons were provided. The patient had to categorize human faces (button 1) versus other stimuli (face-like and matched controls; button 2) appearing at the center of the display by a button press at stimulus offset (after 1000ms of stimulus presentation). Faces appeared in 1 out of 3 trials. Stimulus size measured 8°. Only correct trials were included for analysis. We recorded on day 4 (277 correct trials) and on day 6 (227 correct trials) after array implantation and pooled the data over these two sessions.

### Neurophysiology data analysis

All data analysis was performed using custom-written Matlab (the MathWorks, MA, USA) scripts.

#### SUA/MUA analysis

Net average spike rate was calculated by subtracting the baseline spike rate (0-200 ms before stimulus onset) in each trial. Next, the spike rate per trial was normalized by dividing by the average maximum response calculated in the [25-250] ms interval after stimulus onset. The significance of visual responses and of response differences between conditions was assessed by permutation tests, in which real data were randomly distributed over different conditions 1000 times. The differences between two conditions were calculated for every permutation, and compared with the actual difference between conditions. The signal to noise ratio of the example neuron was calculated by dividing the valley-to-peak with the width of the band before the valley. We calculated d’ values for the contrast faces vs objects and faces vs bodies. Latencies were determined as the first of 2 consecutive 20 ms bins with a spike rate higher than the average baseline plus 2 times the standard error. Selectivity latencies were determined as the first of 2 consecutive 20 ms bins in which the spike rate in one condition exceeded the average spike rate (plus 2 times the standard error) in the other condition. We analyzed the neuronal selectivity by ranking the average spike rate for each individual face-like stimulus and applying the same ranking to the face-like match stimuli. Responses to face stimuli were ranked as well. The 95% confidence interval (CI) for the regression lines are shown in Table S1, those of the matched control did not statistically different from zero.

#### LFP analysis

For every trial, the time-frequency power spectrum was calculated using Morlet’s wavelet analysis techniques, with spectro-temporal resolution equal to 7, after filtering with a 50 Hz notch filter (FieldTrip Toolbox, Donders Institute, The Netherlands). To remove filter artifacts at the beginning and end of the trial, the first and last 100 ms of each trial were discarded. Power was normalized per trial by dividing the power trace per frequency by the average power for this frequency in the 100 ms interval before stimulus onset. To exclude trials containing possible artifacts in the LFP recordings, maximum values of the continuous LFP signal were determined and trials with maximum values above the 95th percentile were removed. We analyzed the LFP power in high frequency bands (high-gamma between 80 and 120 Hz). All statistics on LFP data were obtained using permutation tests, where real data were randomly distributed over all different conditions 1000 times. The differences between two conditions were calculated for every permutation, and compared with the actual difference between conditions. The latency of the high-gamma response was defined as the first of five consecutive timestamps (in ms) in which the average power minus 2 standard errors was higher than 1 (= average power of the normalized baseline). The high-gamma latency for selectivity between conditions was defined as the first of two consecutive samples in which the average power for condition a minus 2 standard errors was higher than the average power for condition b.

#### RF mapping

The average SUA and high-gamma power were calculated during stimulus presentation for each stimulus position and filtered with a Gaussian (sigma: 0.5). To calculate RF size, we constructed RF maps by interpolating the neuronal responses between all positions tested across the 50 × 30 degree display area and then calculating the number of pixels in the RF map with a response higher than 50% of the maximum response.

## Acknowledgements

We thank Stijn Verstraeten, Piet Kayenbergh, Gerrit Meulemans, Marc De Paep, Wouter Depuydt, Inez Puttemans, Christophe Ulens, Anaïs Van Hoylandt, Ron Peeters and Evy Cleeren for technical assistance. We thank Astrid Hermans and Sara De Pril for administrative support.

## Data sharing plan

All data underlying this study will be shared on the Dryad server. Restriction of data availability may apply according to European Union General Data Protection Regulation (GDPR).

## Supplementary information

**Table S1.** The 95% CI for the regression lines for ranking face-like objects and faces. When applying the face-like ranking to the matched control objects, the 95% CI for the regression lines did not statistically differ from zero. Asterisk indicates significance at p<0.05

|  | ranked face-like | matched control | ranked faces |
| --- | --- | --- | --- |
| SUA | [0.0187, 0.00231]* | [-0.0016, 0.0012] | [0.0211, 0.0258]* |
| MUA | [0.0238, 0.00275]* | [-0.0015, 0.0012] | [0.0227, 0.0262]* |
| LFP | [0.0836, 0.1092]* | [-0.0040, 0.0045] | [0.1232, 0.1676]* |

**Figure S1.**
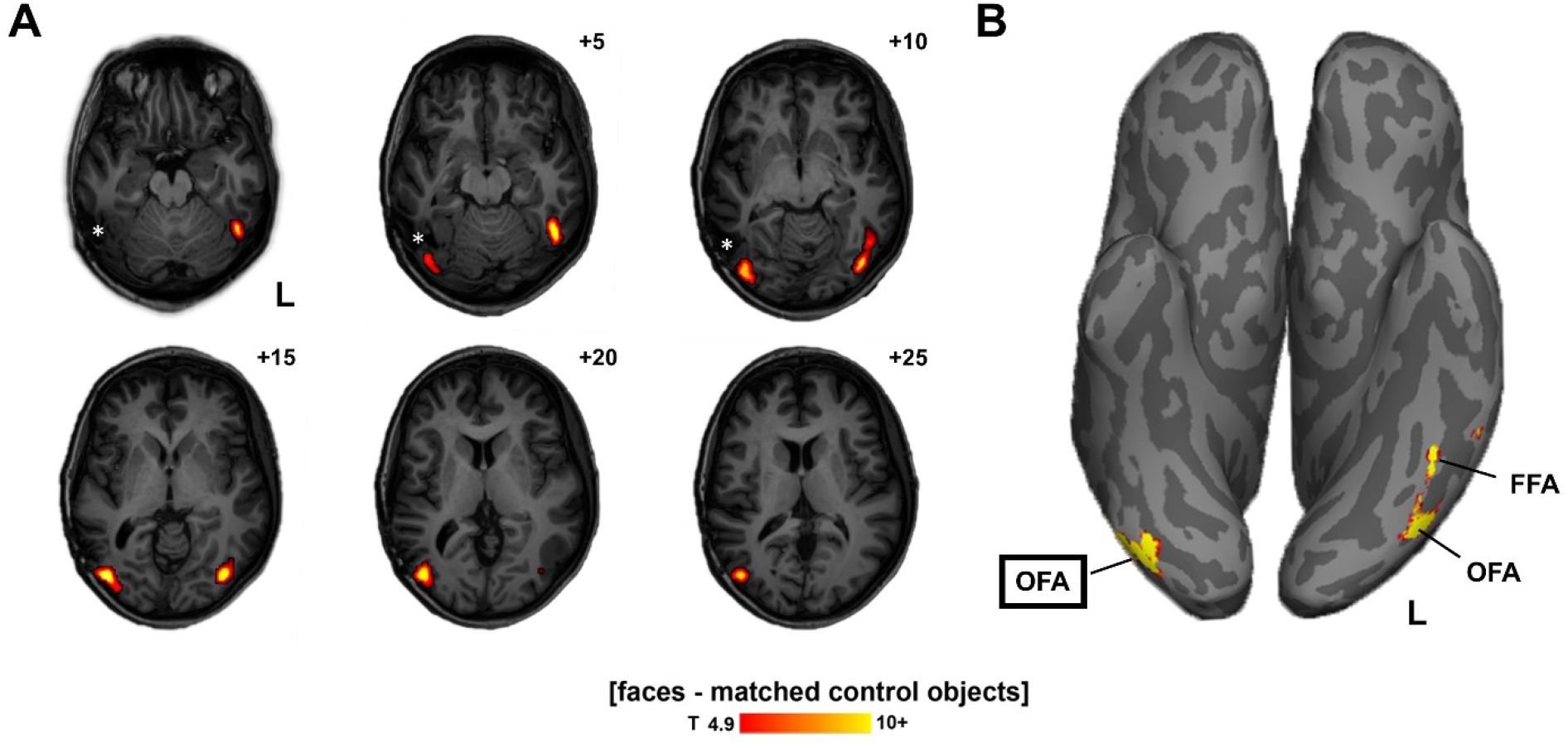
Face-selective regions (all activations for contrast faces>objects; t>4.9) (A) Activations shown on T1 weighted imaging in patient space. Slice distance is indicated in mm, referenced to the most caudal activation on the left side. The lesion after previous surgery is marked with an asterisk. (B) Ventral view on the inflated brain. The OFA was bilaterally present, the FFA only on the left side.

**Figure S2.**
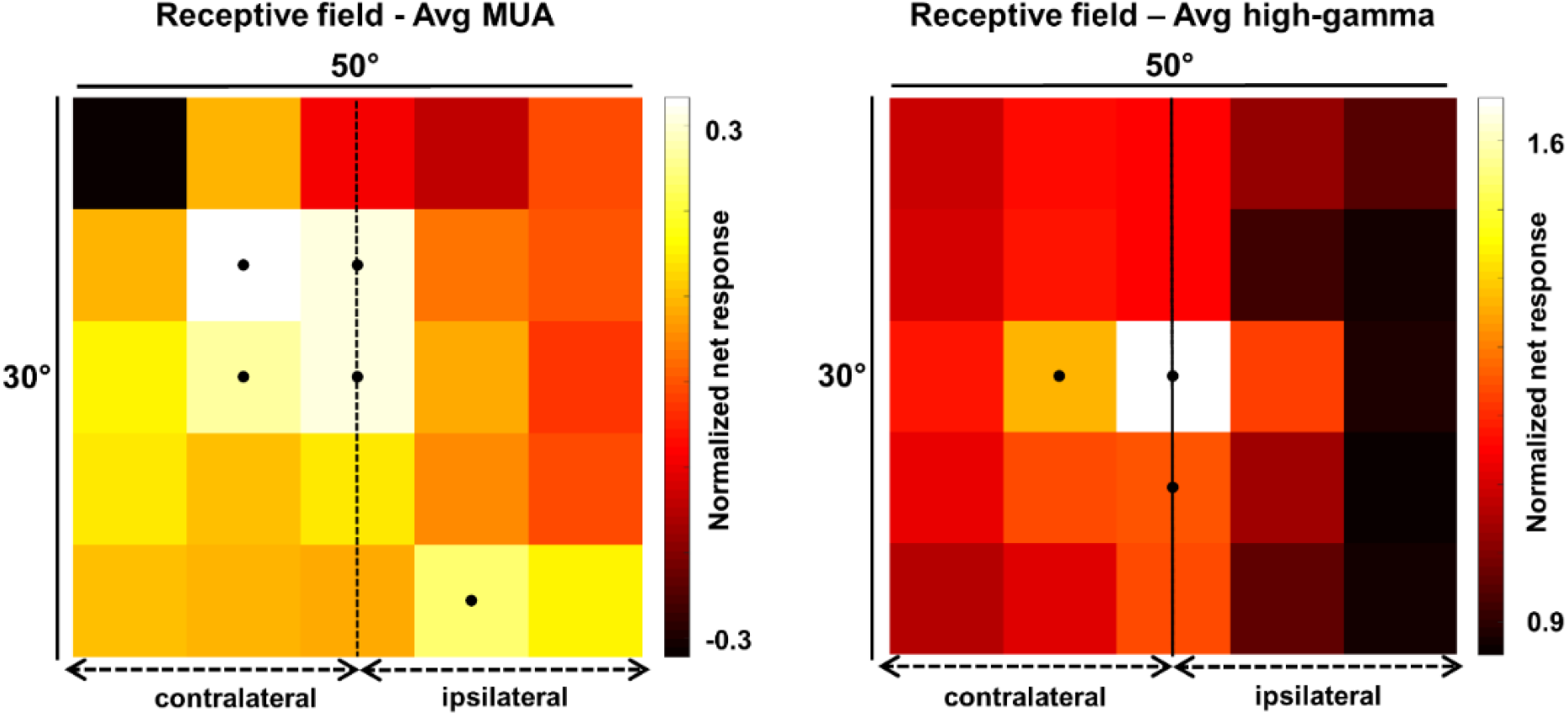
Average receptive field maps. MUA, areas with >50% max response are indicated with a black dot. High-gamma LFP, areas with >75% max response are indicated with a black dot. For both average MUA and high-gamma a preference for the contralateral hemifield exists.

**Figure S3.**
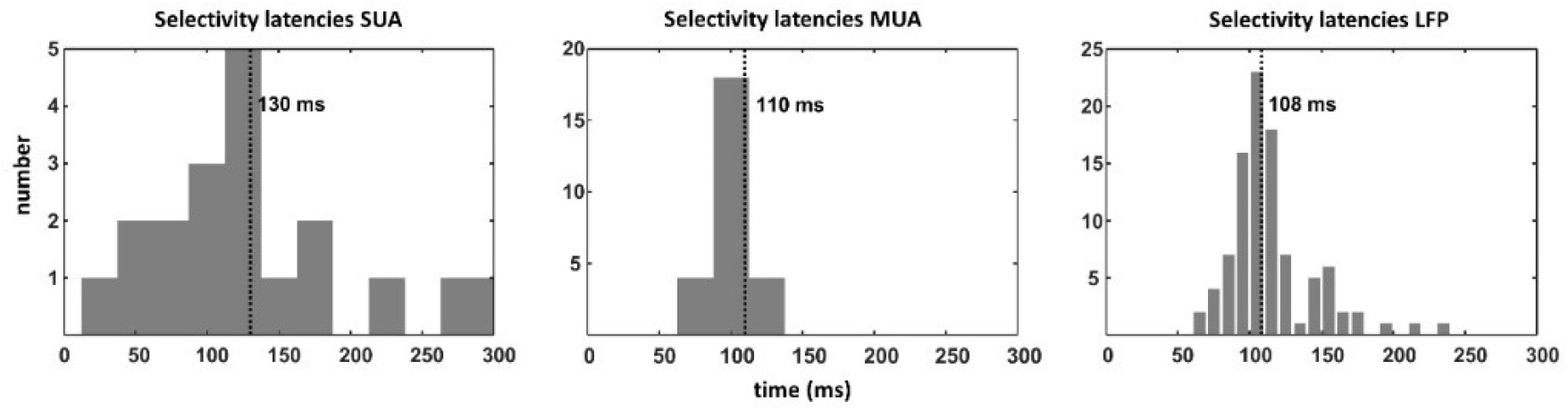
Distribution of selectivity latencies for faces > objects. The number of cells (SUA) or electrodes (MUA/LFP) are depicted on the Y-axis. Median selectivity latencies are marked with dashed lines. (Left) Single-unit activity (SUA). (Middle) Multi-unit activity (MUA). (Right) High-gamma LFP.

**Figure S4.**
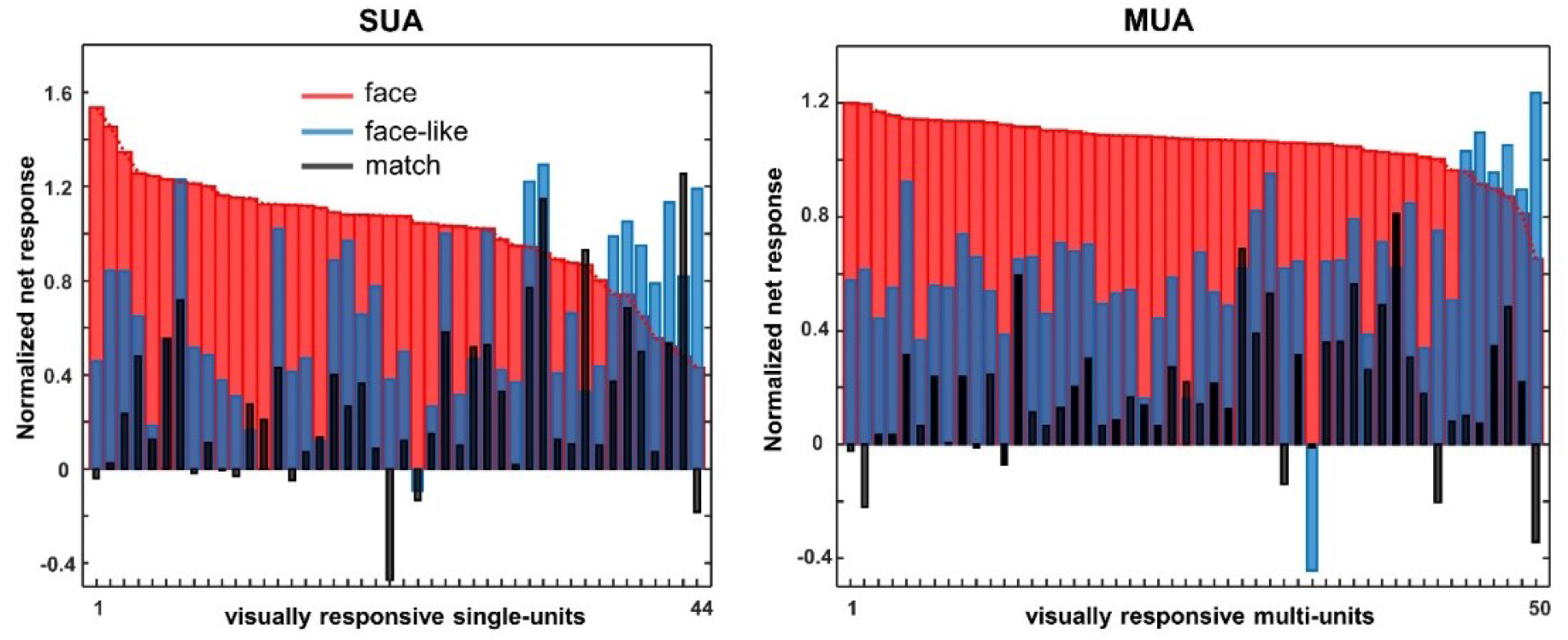
Face pareidolia visually responsive units. Average responses to faces (red), face-like objects (blue) and matched control objects (black) for all visually responsive single units (N=44) and multi-units (N=50).

**Figure S5.**
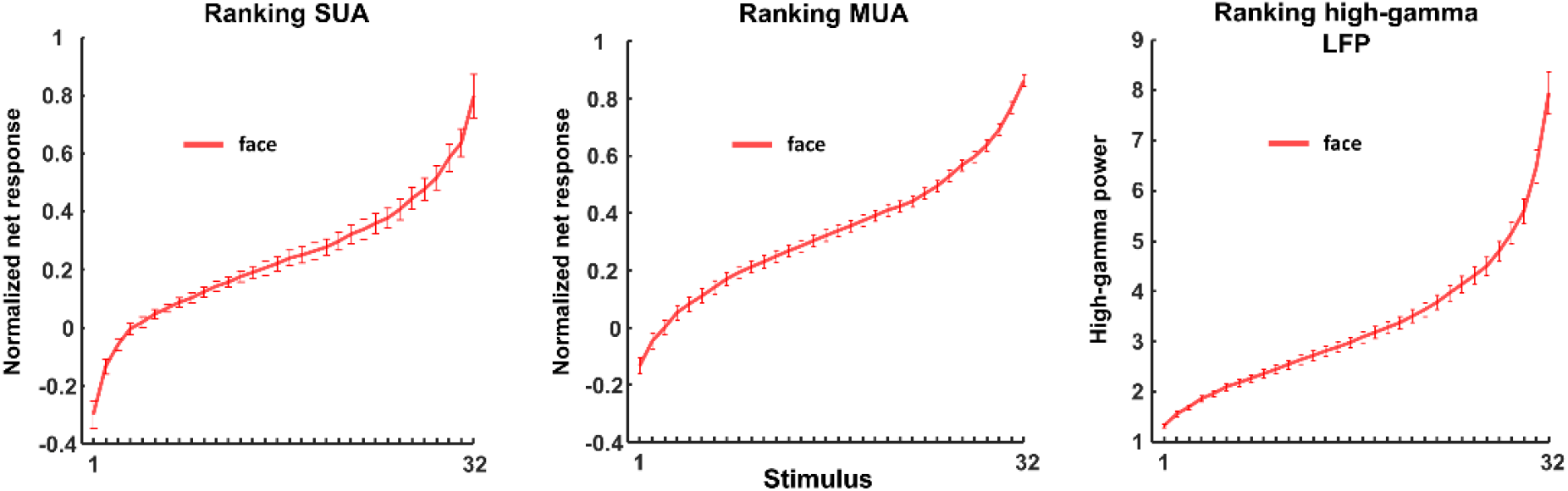
Average ranking for the 32 face stimuli of the pareidolia experiment. (Left) Single-unit activity (SUA). (Center) Multi-unit activity (MUA). (Right) High-gamma LFP. The selectivity for faces was considerable; only 22% (for SUA), 31% (for MUA) and 25% (for high-gamma) of stimuli evoked responses higher than 50% of the maximum response.

## References

Afraz, A., Boyden, E. S., & DiCarlo, J. J. (2015). Optogenetic and pharmacological suppression of spatial clusters of face neurons reveal their causal role in face gender discrimination. Proc Natl Acad Sci U S A, 112(21), 6730–6735. doi: 10.1073/pnas.1423328112

Afraz, S. R., Kiani, R., & Esteky, H. (2006). Microstimulation of inferotemporal cortex influences face categorization. Nature, 442(7103), 692–695. doi: 10.1038/nature04982

Ajiboye, A. B., Willett, F. R., Young, D. R., Memberg, W. D., Murphy, B. A., Miller, J. P., … Kirsch, R. F. (2017). Restoration of reaching and grasping movements through brain-controlled muscle stimulation in a person with tetraplegia: a proof-of-concept demonstration. Lancet, 389(10081), 1821–1830. doi: 10.1016/S0140-6736(17)30601-3

Allison, T., Puce, A., Spencer, D. D., & McCarthy, G. (1999). Electrophysiological studies of human face perception. I: Potentials generated in occipitotemporal cortex by face and non-face stimuli. Cereb Cortex, 9(5), 415-430.

Aparicio, P. L., Issa, E. B., & DiCarlo, J. J. (2016). Neurophysiological Organization of the Middle Face Patch in Macaque Inferior Temporal Cortex. J Neurosci, 36(50), 12729–12745. doi: 10.1523/JNEUROSCI.0237-16.2016

Axelrod, V., Rozier, C., Malkinson, T. S., Lehongre, K., Adam, C., Lambrecq, V., … Naccache, L. (2019). Face-selective neurons in the vicinity of the human fusiform face area. Neurology, 92(4), 197–198. doi: 10.1212/WNL.0000000000006806

Bell, A. H., Malecek, N. J., Morin, E. L., Hadj-Bouziane, F., Tootell, R. B., & Ungerleider, L. G. (2011). Relationship between functional magnetic resonance imaging-identified regions and neuronal category selectivity. J Neurosci, 31(34), 12229–12240. doi: 10.1523/JNEUROSCI.5865-10.2011

Bouvier, S. E., & Engel, S. A. (2006). Behavioral deficits and cortical damage loci in cerebral achromatopsia. Cereb Cortex, 16(2), 183–191. doi: 10.1093/cercor/bhi096

Chang, L., & Tsao, D. Y. (2017). The Code for Facial Identity in the Primate Brain. Cell, 169(6), 1013–1028 e1014. doi: 10.1016/j.cell.2017.05.011

Collinger, J. L., Wodlinger, B., Downey, J. E., Wang, W., Tyler-Kabara, E. C., Weber, D. J., … Schwartz, A. B. (2013). High-performance neuroprosthetic control by an individual with tetraplegia. Lancet, 381(9866), 557–564. doi: 10.1016/S0140-6736(12)61816-9

Davidesco, I., Zion-Golumbic, E., Bickel, S., Harel, M., Groppe, D. M., Keller, C. J., … Malach, R. (2014). Exemplar selectivity reflects perceptual similarities in the human fusiform cortex. Cereb Cortex, 24(7), 1879–1893. doi: 10.1093/cercor/bht038

Davis, T. S., Wark, H. A., Hutchinson, D. T., Warren, D. J., O’Neill, K., Scheinblum, T., … Greger, B. (2016). Restoring motor control and sensory feedback in people with upper extremity amputations using arrays of 96 microelectrodes implanted in the median and ulnar nerves. J Neural Eng, 13(3), 036001. doi: 10.1088/1741-2560/13/3/036001

Decramer, T., Premereur, E., Uytterhoeven, M., Van Paesschen, W., van Loon, J., Janssen, P., & Theys, T. (2019). Single-cell selectivity and functional architecture of human lateral occipital complex. PLoS Biol, 17(9), e3000280. doi: 10.1371/journal.pbio.3000280

Fernandez, E., Greger, B., House, P. A., Aranda, I., Botella, C., Albisua, J., … Normann, R. A. (2014). Acute human brain responses to intracortical microelectrode arrays: challenges and future prospects. Front Neuroeng, 7, 24. doi: 10.3389/fneng.2014.00024

Flesher, S. N., Collinger, J. L., Foldes, S. T., Weiss, J. M., Downey, J. E., Tyler-Kabara, E. C., … Gaunt, R. A. (2016). Intracortical microstimulation of human somatosensory cortex. Sci Transl Med, 8(361), 361ra141. doi: 10.1126/scitranslmed.aaf8083

Fried, I., MacDonald, K. A., & Wilson, C. L. (1997). Single neuron activity in human hippocampus and amygdala during recognition of faces and objects. Neuron, 18(5), 753–765.

Gauthier, I., Skudlarski, P., Gore, J. C., & Anderson, A. W. (2000). Expertise for cars and birds recruits brain areas involved in face recognition. Nat Neurosci, 3(2), 191–197. doi: 10.1038/72140

Grill-Spector, K., Knouf, N., & Kanwisher, N. (2004). The fusiform face area subserves face perception, not generic within-category identification. Nat Neurosci, 7(5), 555–562. doi: 10.1038/nn1224

Grill-Spector, K., Weiner, K. S., Kay, K., & Gomez, J. (2017). The Functional Neuroanatomy of Human Face Perception. Annu Rev Vis Sci, 3, 167–196. doi: 10.1146/annurev-vision-102016-061214

Haxby, J. V., Hoffman, E. A., & Gobbini, M. I. (2000). The distributed human neural system for face perception. Trends Cogn Sci, 4(6), 223–233. doi: 10.1016/s1364-6613(00)01482-0

Hemond, C. C., Kanwisher, N. G., & Op de Beeck, H. P. (2007). A preference for contralateral stimuli in human object- and face-selective cortex. PLoS One, 2(6), e574. doi: 10.1371/journal.pone.0000574

Hochberg, L. R., Bacher, D., Jarosiewicz, B., Masse, N. Y., Simeral, J. D., Vogel, J., … Donoghue, J. P. (2012). Reach and grasp by people with tetraplegia using a neurally controlled robotic arm. Nature, 485(7398), 372–375. doi: 10.1038/nature11076

Hochberg, L. R., Serruya, M. D., Friehs, G. M., Mukand, J. A., Saleh, M., Caplan, A. H., … Donoghue, J. P. (2006). Neuronal ensemble control of prosthetic devices by a human with tetraplegia. Nature, 442(7099), 164–171. doi: 10.1038/nature04970

House, P. A., MacDonald, J. D., Tresco, P. A., & Normann, R. A. (2006). Acute microelectrode array implantation into human neocortex: preliminary technique and histological considerations. Neurosurg Focus, 20(5), E4.

Issa, E. B., & DiCarlo, J. J. (2012). Precedence of the eye region in neural processing of faces. J Neurosci, 32(47), 16666–16682. doi: 10.1523/JNEUROSCI.2391-12.2012

Jacques, C., Witthoft, N., Weiner, K. S., Foster, B. L., Rangarajan, V., Hermes, D., … Grill-Spector, K. (2016). Corresponding ECoG and fMRI category-selective signals in human ventral temporal cortex. Neuropsychologia, 83, 14–28. doi: 10.1016/j.neuropsychologia.2015.07.024

Jonas, J., Descoins, M., Koessler, L., Colnat-Coulbois, S., Sauvee, M., Guye, M., … Maillard, L. (2012). Focal electrical intracerebral stimulation of a face-sensitive area causes transient prosopagnosia. Neuroscience, 222, 281–288. doi: 10.1016/j.neuroscience.2012.07.021

Jonas, J., Jacques, C., Liu-Shuang, J., Brissart, H., Colnat-Coulbois, S., Maillard, L., & Rossion, B. (2016). A face-selective ventral occipito-temporal map of the human brain with intracerebral potentials. Proc Natl Acad Sci U S A, 113(28), E4088–4097. doi: 10.1073/pnas.1522033113

Kanwisher, N., McDermott, J., & Chun, M. M. (1997). The fusiform face area: a module in human extrastriate cortex specialized for face perception. J Neurosci, 17(11), 4302–4311.

Kreiman, G., Koch, C., & Fried, I. (2000). Imagery neurons in the human brain. Nature, 408(6810), 357–361. doi: 10.1038/35042575

Liu, H., Agam, Y., Madsen, J. R., & Kreiman, G. (2009). Timing, timing, timing: fast decoding of object information from intracranial field potentials in human visual cortex. Neuron, 62(2), 281–290. doi: 10.1016/j.neuron.2009.02.025

Liu, J., Li, J., Feng, L., Li, L., Tian, J., & Lee, K. (2014). Seeing Jesus in toast: neural and behavioral correlates of face pareidolia. Cortex, 53, 60–77. doi: 10.1016/j.cortex.2014.01.013

Martinet, L. E., Fiddyment, G., Madsen, J. R., Eskandar, E. N., Truccolo, W., Eden, U. T., … Kramer, M. A. (2017). Human seizures couple across spatial scales through travelling wave dynamics. Nat Commun, 8, 14896. doi: 10.1038/ncomms14896

McCarthy, G., Puce, A., Belger, A., & Allison, T. (1999). Electrophysiological studies of human face perception. II: Response properties of face-specific potentials generated in occipitotemporal cortex. Cereb Cortex, 9(5), 431–444. doi: 10.1093/cercor/9.5.431

McCarthy, G., Puce, A., Gore, J. C., & Allison, T. (1997). Face-specific processing in the human fusiform gyrus. J Cogn Neurosci, 9(5), 605–610. doi: 10.1162/jocn.1997.9.5.605

Nihei, Y., Minami, T., & Nakauchi, S. (2018). Brain Activity Related to the Judgment of Face-Likeness: Correlation between EEG and Face-Like Evaluation. Front Hum Neurosci, 12, 56. doi: 10.3389/fnhum.2018.00056

Parvizi, J., Jacques, C., Foster, B. L., Witthoft, N., Rangarajan, V., Weiner, K. S., & Grill-Spector, K. (2012). Electrical stimulation of human fusiform face-selective regions distorts face perception. J Neurosci, 32(43), 14915–14920. doi: 10.1523/JNEUROSCI.2609-12.2012

Pitcher, D., Charles, L., Devlin, J. T., Walsh, V., & Duchaine, B. (2009). Triple dissociation of faces, bodies, and objects in extrastriate cortex. Curr Biol, 19(4), 319–324. doi: 10.1016/j.cub.2009.01.007

Pitcher, D., Walsh, V., & Duchaine, B. (2011). The role of the occipital face area in the cortical face perception network. Exp Brain Res, 209(4), 481–493. doi: 10.1007/s00221-011-2579-1

Pitcher, D., Walsh, V., Yovel, G., & Duchaine, B. (2007). TMS evidence for the involvement of the right occipital face area in early face processing. Curr Biol, 17(18), 1568–1573. doi: 10.1016/j.cub.2007.07.063

Popivanov, I. D., Jastorff, J., Vanduffel, W., & Vogels, R. (2014). Heterogeneous single-unit selectivity in an fMRI-defined body-selective patch. J Neurosci, 34(1), 95–111. doi: 10.1523/JNEUROSCI.2748-13.2014

Puce, A., Allison, T., & McCarthy, G. (1999). Electrophysiological studies of human face perception. III: Effects of top-down processing on face-specific potentials. Cereb Cortex, 9(5), 445–458. doi: 10.1093/cercor/9.5.445

Quian Quiroga, R., Kraskov, A., Koch, C., & Fried, I. (2009). Explicit encoding of multimodal percepts by single neurons in the human brain. Curr Biol, 19(15), 1308–1313. doi: 10.1016/j.cub.2009.06.060

Quiroga, R. Q., Reddy, L., Kreiman, G., Koch, C., & Fried, I. (2005). Invariant visual representation by single neurons in the human brain. Nature, 435(7045), 1102–1107. doi: 10.1038/nature03687

Rainer, G., Asaad, W. F., & Miller, E. K. (1998). Selective representation of relevant information by neurons in the primate prefrontal cortex. Nature, 393(6685), 577–579. doi: 10.1038/31235

Riley, J. D., Fling, B. W., Cramer, S. C., & Lin, J. J. (2015). Altered organization of face-processing networks in temporal lobe epilepsy. Epilepsia, 56(5), 762–771. doi: 10.1111/epi.12976

Rossion, B. (2008). Constraining the cortical face network by neuroimaging studies of acquired prosopagnosia. Neuroimage, 40(2), 423–426. doi: 10.1016/j.neuroimage.2007.10.047

Rossion, B., & Taubert, J. (2019). What can we learn about human individual face recognition from experimental studies in monkeys? Vision Res, 157, 142–158. doi: 10.1016/j.visres.2018.03.012

Sato, W., Kochiyama, T., Uono, S., Matsuda, K., Usui, K., Inoue, Y., & Toichi, M. (2014). Rapid, high-frequency, and theta-coupled gamma oscillations in the inferior occipital gyrus during face processing. Cortex, 60, 52–68. doi: 10.1016/j.cortex.2014.02.024

Simeral, J. D., Kim, S. P., Black, M. J., Donoghue, J. P., & Hochberg, L. R. (2011). Neural control of cursor trajectory and click by a human with tetraplegia 1000 days after implant of an intracortical microelectrode array. J Neural Eng, 8(2), 025027. doi: 10.1088/1741-2560/8/2/025027

Smith, E. H., Liou, J. Y., Davis, T. S., Merricks, E. M., Kellis, S. S., Weiss, S. A., … Schevon, C. A. (2016). The ictal wavefront is the spatiotemporal source of discharges during spontaneous human seizures. Nat Commun, 7, 11098. doi: 10.1038/ncomms11098

Sorger, B., Goebel, R., Schiltz, C., & Rossion, B. (2007). Understanding the functional neuroanatomy of acquired prosopagnosia. Neuroimage, 35(2), 836–852. doi: 10.1016/j.neuroimage.2006.09.051

Taubert, J., Wardle, S. G., Flessert, M., Leopold, D. A., & Ungerleider, L. G. (2017). Face Pareidolia in the Rhesus Monkey. Curr Biol, 27(16), 2505-2509 e2502. doi: 10.1016/j.cub.2017.06.075

Truccolo, W., Donoghue, J. A., Hochberg, L. R., Eskandar, E. N., Madsen, J. R., Anderson, W. S., … Cash, S. S. (2011). Single-neuron dynamics in human focal epilepsy. Nat Neurosci, 14(5), 635–641. doi: 10.1038/nn.2782

Tsao, D. Y., Freiwald, W. A., Tootell, R. B., & Livingstone, M. S. (2006). A cortical region consisting entirely of face-selective cells. Science, 311(5761), 670–674. doi: 10.1126/science.1119983

Tsao, D. Y., & Livingstone, M. S. (2008). Mechanisms of face perception. Annu Rev Neurosci, 31, 411–437. doi: 10.1146/annurev.neuro.30.051606.094238

Tsao, D. Y., Moeller, S., & Freiwald, W. A. (2008). Comparing face patch systems in macaques and humans. Proc Natl Acad Sci U S A, 105(49), 19514–19519. doi: 10.1073/pnas.0809662105

Wardle, Susan G., Seymour, Kiley, & Taubert, Jessica. (2017). Characterizing the response to face pareidolia in human category-selective visual cortex. bioRxiv. doi: 10.1101/233387

Weiss, S. A., McKhann, G., Jr., Goodman, R., Emerson, R. G., Trevelyan, A., Bikson, M., & Schevon, C. A. (2013). Field effects and ictal synchronization: insights from in homine observations. Front Hum Neurosci, 7, 828. doi: 10.3389/fnhum.2013.00828

